# The evolution of coercive policing in genetically mixed groups: the case of plasmid copy number control

**DOI:** 10.1101/053579

**Authors:** Kyriakos Kentzoglanakis, Sam P. Brown, Richard A. Goldstein

**Affiliations:** Division of Infection & Immunity, University College London, UK; School of Biology, Georgia Institute of Technology, Georgia, USA

## Abstract

Policing is a widespread mechanism regulating cooperation in both human and animal social groups. Policing can promote the evolution and maintenance of cooperation among non-relatives by tying the reproductive success of individuals to the efficiency and success of the group. In this paper, we investigate the evolution of reproductive policing using a multi-scale computational model inspired by the copy number control system of conjugative bacterial plasmids. Our results show that the repression of competition through policing can evolve across a very broad range of migration (plasmid conjugation) rates, improving system-level performance and bringing efficiency gains to the group beyond those achievable by pure self-restraint. Reproductive policing acts to increase genetic relatedness by reducing variation in group size which, in turn, reduces the heterogeneity of the plasmid population. When among-group migration is high, coercive policing strategies are favoured, characterized by high levels of policing coupled with relatively lower obedience. Coercive policing strategies preferentially limit the reproduction of rival lineages while, at the same time, maintaining effective collective reproductive control.

**Author Summary:** The emergence and maintenance of cooperation is a topic of great importance in evolutionary biology. The evolution of cooperation has been explained in the context of kin selection when there is sufficient genetic relatedness among interacting individuals. When there is insufficient relatedness, the presence of alternative mechanisms, such as mutual policing, can promote the evolution and maintenance of cooperation by tying the reproductive success of individual to the efficiency and success of the group. In this paper, we investigate the evolution of reproductive policing using an agent-based computational model inspired by a simple and elegant biological example: replication control among conjugative plasmids, a class of molecular symbionts of bacterial hosts. Our results show that the repression of competition through policing evolves and improves plasmid group performance beyond levels achievable by self-restraint, across a very broad range of migration rates. Under conditions of high migration (frequent conjugation), we observe the evolution of coercive policing strategies that limit the reproduction of rival lineages by investing disproportionately in policing relative to their obedience to the policing trait.

## Introduction

Cooperation is a ubiquitous feature of biology. From microbes to humans, examples of individual self-sacrifice to promote the interests of others are widely observed across the spectrum of biological organization. The widespread presence of helping behaviours presents a puzzle: how are these traits maintained in the presence of “cheats”, individuals that reduce or abandon cooperation in favour of more competitive interactions [1–4]. When individuals preferentially interact with kin, high genetic relatedness can favour cooperation, as help to others promotes the transmission of shared “cooperation” alleles [5,6]. However, kin selection cannot account for the maintenance of costly cooperation among non-relatives. When the costs of cooperation are less than the expected benefits returned by reciprocation of aid, then cooperation can still be favoured due to direct fitness benefits [7,8]. The relative costs of cooperation can be further mitigated by additional mechanisms of enforcement, such as mutual policing [9]. Investments in such mechanisms diminish the rewards of selfish behaviours resulting in a leveling of the playing field within a group through repression of competition, thus tying individual reproductive success to the success of the group.

Frank [10] developed a simple and influential model outlining the logic of policing investments in social groups. Under this approach (see also [9,11,12]), policing is redundant in the absence of within-group genetic conflict (high relatedness), as cooperation is predicted to be effectively maintained purely via kin selection. In contrast to these findings, we recently presented a model of policing (inspired by the collective control of reproduction among plasmids) where policing could evolve under a high relatedness scenario, due to efficiency gains improving group productivity [13].

The replication control system of plasmids constitutes a simple and elegant example of a policing mechanism. Plasmids are molecular symbionts of bacteria, extra-chromosomal genetic elements that replicate autonomously within bacterial cells. Intra-cellular replication is limited by the collective production of plasmid-coded replication inhibitor molecules (the policing trait), which bind to and inhibit the replication of individual plasmids. Individual plasmid investments in the shared inhibitor molecule generate an effective negative-feedback mechanism controlling within-cell plasmid copy number within narrow bounds [15–17]. Kentzoglanakis et al. [13] decomposed this mechanism of plasmid copy number control (CNC) into three distinct plasmid traits, namely “policing” (inhibitor molecule production), “obedience” (affinity for inhibitor binding) and “intrinsic selfishness” (maximal replication rate), which can co-evolve to improve the collective behaviour of plasmids within host cells, thus reducing the risks of over-and under-exploitation of their shared cellular resource.

The model of Kentzoglanakis et al. [13] demonstrated clear efficiency benefits from the evolution of a policing mechanism even in a high relatedness scenario. Specifically, the plasmids were limited to purely vertical (mother to daughter cell) transmission, ensuring that within-cell genetic variation was minimized relative to genetic variation among cells. However, plasmids are also capable of highly variable degrees of autonomous horizontal transmission [18–21]. The introduction of between-group migration will reduce relatedness among interacting individuals and, therefore, increase the potential benefits arising from the enforcement of cooperation.

In this paper, we investigate the evolution and the effect of policing on hosts and plasmids across a broad range of mixing regimes. Our general strategy is to use an in-silico agent-based model of plasmid evolution with and without an explicit CNC (policing) mechanism, and under varying degrees of plasmid migration, which implies a variable relatedness. Specifically, we ask whether the policing mechanism evolves and, if so, whether it improves collective performance above pure kin-selected restraint across a range of relatedness regimes. In addition, we investigate the effects of policing on the population's genetic structure and we decompose the selective forces acting on the plasmid replication parameters using the Price equation in order to elucidate the effects of migration on the evolution of policing in the context of plasmid replication control.

## Models

As a baseline, we use a previously described model [13,14] specifying the mechanics of the asynchronous growth, division and death of hosts in a bacterial population, as well as the independent replication of plasmids within hosts. We extend this model here with provisions for the migration of plasmids among hosts. We consider the individual growth of each host *i* in a population where growth is represented by a biomass variable Ω_i_ which is updated at each time step of the simulation according to:

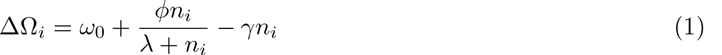

where *n_i_* is the total number of plasmid copies in the host and Ω_0_ is the cell’s basal growth rate. Parameters and A characterize the beneficial contribution of the plasmid trait to cell metabolism which saturates at large *n*, while 7 defines the cost of plasmid maintenance (including the costs of gene expression, replication etc.) which increases in proportion to the copy number [22–24]. When Ω_*i*_≥2Ω_0_ (with Ω_0_ = 1), the cell divides and its plasmids are segregated binomially among the two daughter cells, while when Ω_*i*_<0 the cell dies as a result of a negative balance between metabolic benefits and costs. Equation 1 yields a copy number 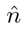 that maximizes the host’s growth rate ΔΩ given by:

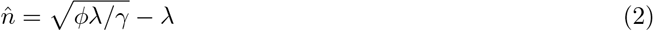

Within each host *i* in the population, plasmids replicate independently by competing in the intracellular replication pool. This competition is subject to regulation by the plasmid-coded CNC system, on the basis of the production of plasmid-coded trans-acting replication inhibitors that bind to and deactivate a plasmid-specific target, thereby down-regulating the individual plasmid replication rate [25]. The replication rate of plasmid *j* in host *i* is given by:

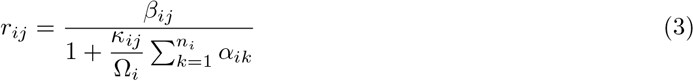

where β is the plasmid’s basal replication rate, and *κ_ij_* is its binding affinity to the generic replication inhibitor, which is synthesized by each plasmid at a rate *α_ij_/τ*, where τ is the (short) average lifetime of the inhibitor. The replication of a plasmid includes the possibility of mutation with probability μ, in which case exactly one of the parameter values (*β_ij_*, *κ_ij_* or *α_ij_*chosen at random) is slightly modified.

We also define migration among groups as the probability *p_c_* that an infected host (donor) will successfully transmit a single copy of a plasmid to another randomly selected host (recipient) at each time step. The probability *p_c_* encapsulates various factors that determine the frequency of cell-cell contacts, such as population density, and the likelihood of a successful horizontal transmission event [26],such as the existence and efficiency of surface/entry exclusion mechanisms [27] etc. We assume that the population of hosts is well-mixed and that each plasmid in a donor host is equally probable to be transmitted to the recipient.

## Results

In our stochastic simulations, we are interested in the evolution of plasmid replication control across a wide range of migration rates. In particular, we consider two distinct scenarios for our simulations: in the first one, plasmids replicate solely on the basis of self-restraint (NO-CNC; β evolves, while κ = *α* = 0), while, in the second one, the replication of plasmids is subject to collective restraint through policing (CNC; β, κ and *α* all evolve). In the latter case, the simulations start with an inactive CNC mechanism, whereby initially plasmids neither produce replication inhibitors (*α* = 0) nor respond to their presence (κ = 0). In all simulations we set *μ* = 0.005 (mutation probability), ω_0_ = 0.05, ϕ = 0.1, *γ* = 0.003 and λ = 3 (Equation 1), yielding an optimal copy number 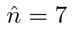 (Equation 2).

### The effects of migration on hosts

We begin by investigating how the migration of plasmids influences the performance of hosts in the population. Figure 1 demonstrates that the evolution of the policing mechanism (CNC) enhances the performance of hosts over a broad range of migration rates compared to the case where policing is absent (NO-CNC) and plasmids replicate on the basis of self-restraint. Specifically, host growth and death rates are significantly improved when plasmid replication is subject to copy number control. The system’s behaviour in response to increasing migration is generally characterized by the growth of copy numbers in the population. In the CNC case, this corresponds to higher metabolic loads that impose a detrimental effect on host growth, as captured by the decreasing average host growth rates and the increasing average death rates, especially at high levels of migration (*p_c_* ≥0.02). For *p_c_* > 0.1, the extremely high plasmid turnover mediates the collapse of plasmid obedience to policing and the subsequent extinction of the population as a result of excessive plasmid replication.

**Figure 1.**
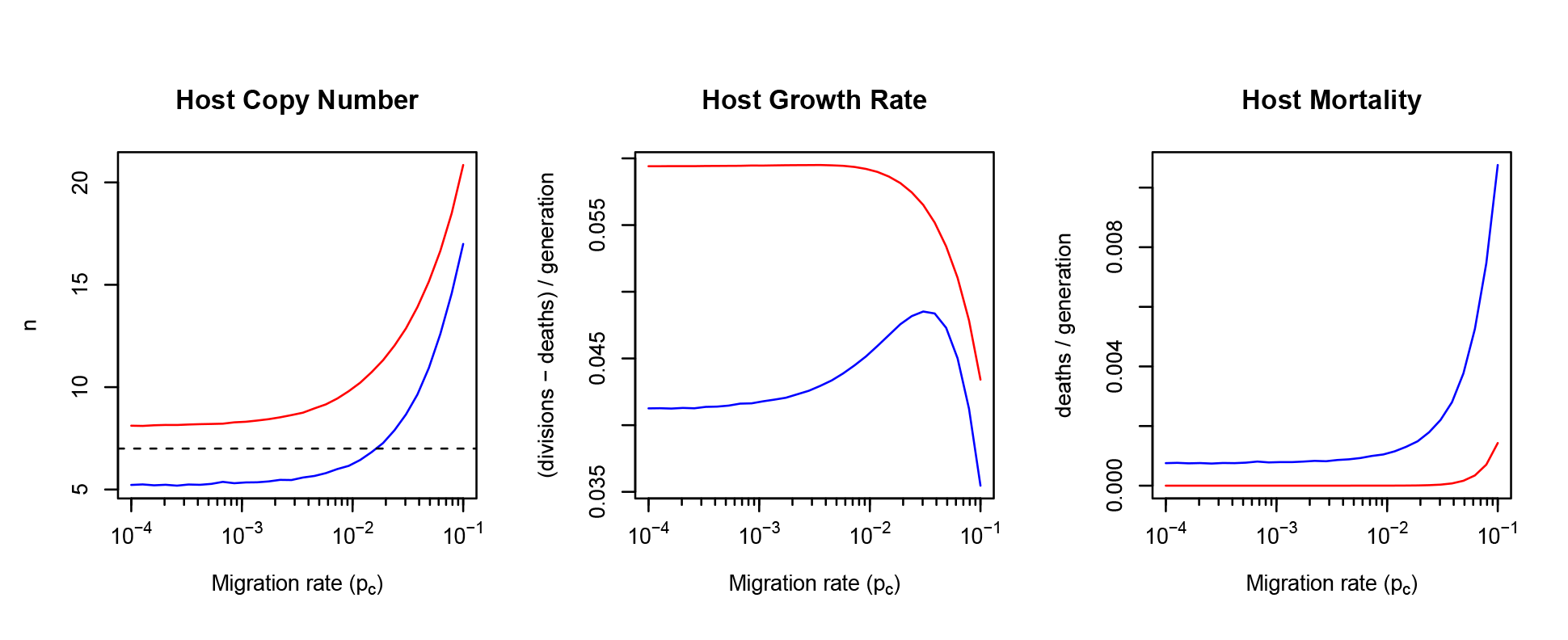
Policing enhances collective performance over a broad range of migration rates. The mean per-host copy number (left) and the mean host growth (middle) and death (right) rates in the presence (CNC - red) and the absence (NO-CNC - blue) of policing, calculated from the equilibrium phase (second half) of the simulation dynamics and averaged across 100 independent simulations for each value of the migration probability *p_c_*. The horizontal dashed lin in the leftmost panel denotes the copy number 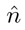 that is optimal for host growth (Equation 2).

The fact that average host copy numbers are lower in the NO-CNC case (Figure 1, left), when intuition dictates that the presence of policing would regulate copy numbers more efficiently, can be explained by the inherent instabilities of intra-cellular plasmid replication in the absence of policing. These instabilities are characterized by the consistent under-or over-replication of plasmids in the absence of copy number control [13]. The resulting copy number distributions, shown in Figure 2 (NO-CNC), demonstrate that, as migration become more frequent, the system undergoes a transition from the regime of under-replication (low migration) to that of over-replication (high migration) crossing the point of copy number optimality in which the mean copy number is equal to the optimal copy number 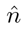 (Equation 2). This effect is reflected in the concavity of the NO-CNC host growth curve (Figure 1, middle), that indicates the consistent under-replication of plasmids before the peak and their consistent over-replication beyond the peak. Even at this peak, however, the growth rate is lower than with copy number control, due to a less favorable distribution of copy numbers (Figure 2).

**Figure 2.**
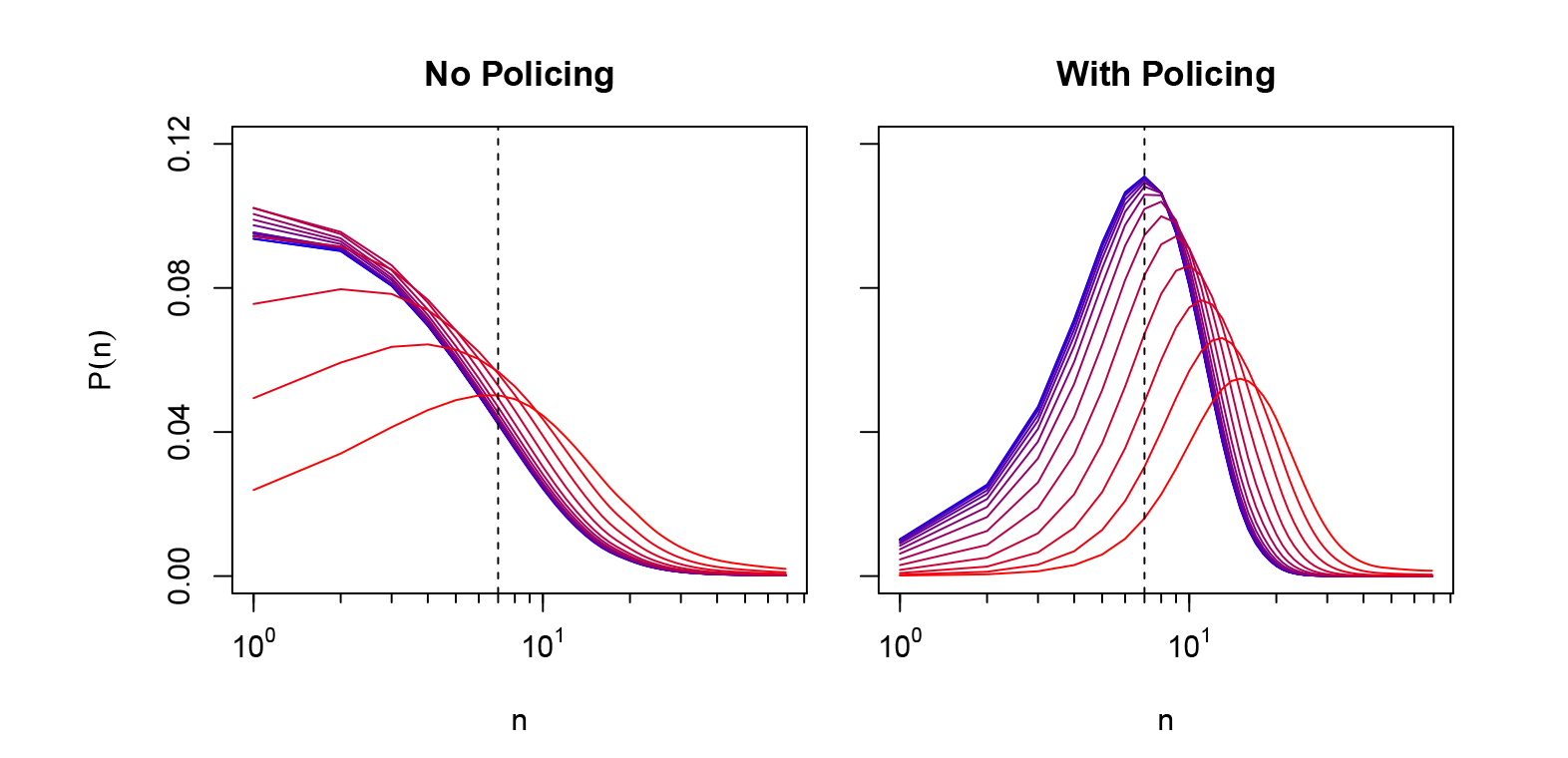
The effects of migration on copy number distributions. The distributions of plasmidcopy numbers in the population, calculated by aggregating the results of 100 independent simulations, from low (*p_c_* = 10^−4^ − blue) to high (*p_c_* = 10^−1^ − red) migration rates. The vertical dashed line denotes the copy number 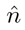 that is optimal for host growth (Equation 2).

### The effects of migration on plasmids

The effects of migration on plasmid dynamics are dominated by the growth of copy numbers in the population (Figure 1, left). Figure 3 (top left) demonstrates that, as migration becomes more frequent and host copy numbers grow, the per-plasmid replication rate increases in the absence of policing (NO-CNC), indicating a deterioration of reproductive control among plasmids due to increasing competition in the intra-cellular replication pool. On the contrary, when the policing mechanism is functional (CNC), the per-plasmid replication rate declines as the accumulation of replication inhibitors (in proportion to the growing host copy number) effectively down-regulates intra-cellular plasmid replication.

**Figure 3.**
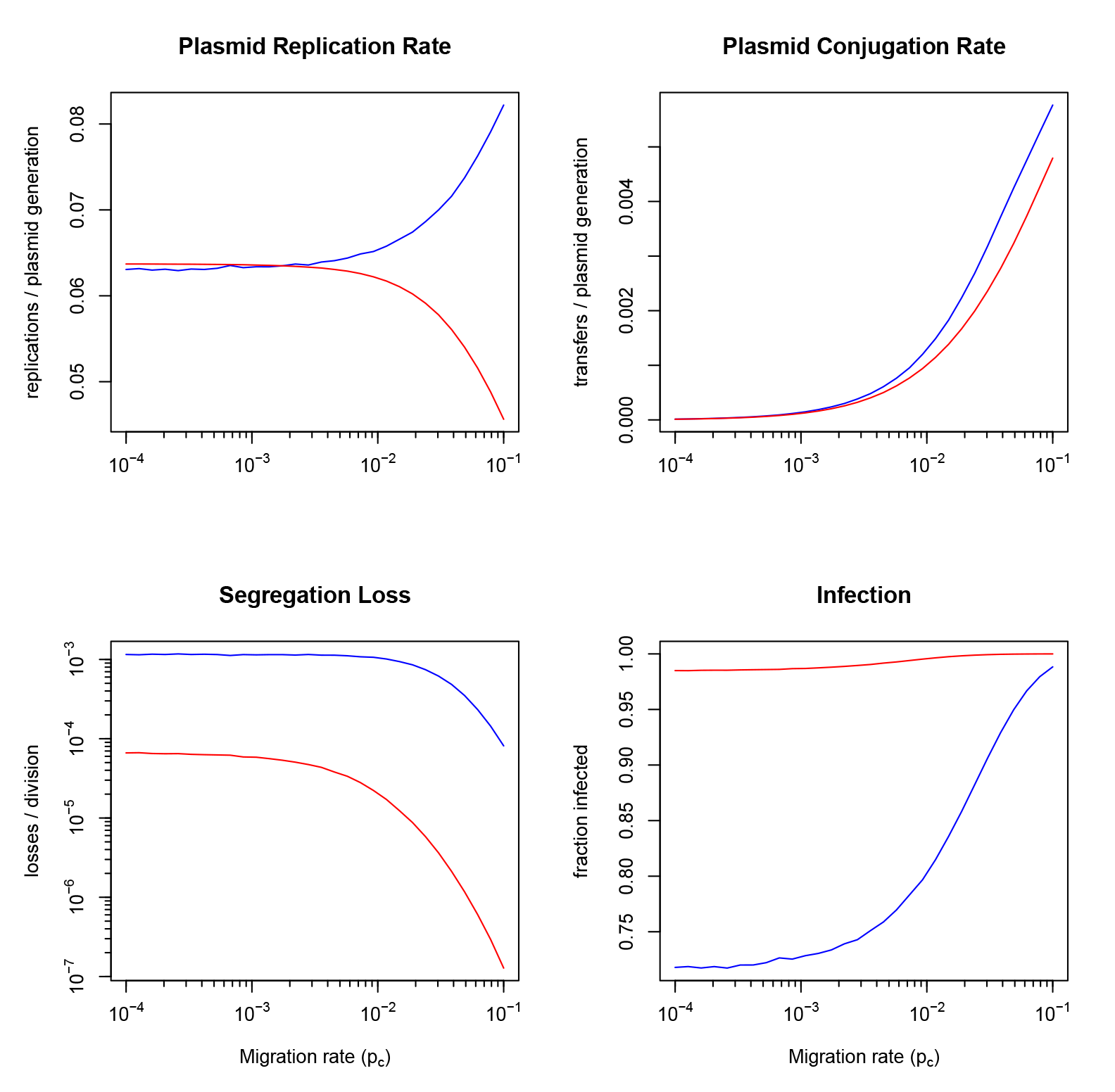
In the absence of policing, reproductive control weakens as migration increases. Mean per-plasmid replication (top left) and conjugation (top right) rates as well as the probabilities of segregation loss (bottom left) and host infection (bottom right) in the presence (CNC - red) and the absence (NO-CNC - blue) of policing, calculated from the equilibrium phase (second half) of the simulation dynamics and averaged across 100 independent simulations for each value of the migration probability *p_c_*.

Increasing migration rates and copy number growth also result in the increase of the per-plasmid conjugation rate (Figure 3 - top right) as well as the progressive minimization of the probability of segregational loss (see Figure 3 - bottom left). The latter occurs when a plasmid-free daughter cell results from the division of an infected parent cell. The consequent improvements in the fidelity of vertical plasmid transmission are present in both cases of self-restraint (NO-CNC) and policing (CNC), however in the presence of policing (CNC), the probability of segregational loss is orders of magnitude lower compared to the NO-CNC case, due to increased efficiency of control over the local population size. Furthermore, the presence of policing also promotes the wider spread of plasmids in the population (see Figure 3 - bottom right), as measured by the fraction of plasmid-infected hosts which is significantly larger in the CNC case, across the spectrum of migration rates.

### Policing and relatedness

We proceed by investigating the effects of migration on the equilibrium properties of the policing system itself, as defined by the plasmid parameters κ (obedience) and α (production of policing resources). Figure 4 (left) demonstrates that policing is robust with respect to varying levels *p_c_* of plasmid migration. Specifically, the average obedience 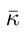 of plasmids to policing, while decreasing slightly, remains at high levels (above 0.9) across the pc spectrum, while the average production 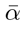 of inhibitors gradually increases until it eventually reaches and maintains its maximum value at 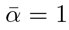. The growth of the average basal replication rate 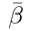 as a function of *p_c_* reflects the intensification of competition among plasmids in the intra-cellular replication pool, which is fuelled by the associated growth of copy numbers in the population (Figure 1).

**Figure 4.**
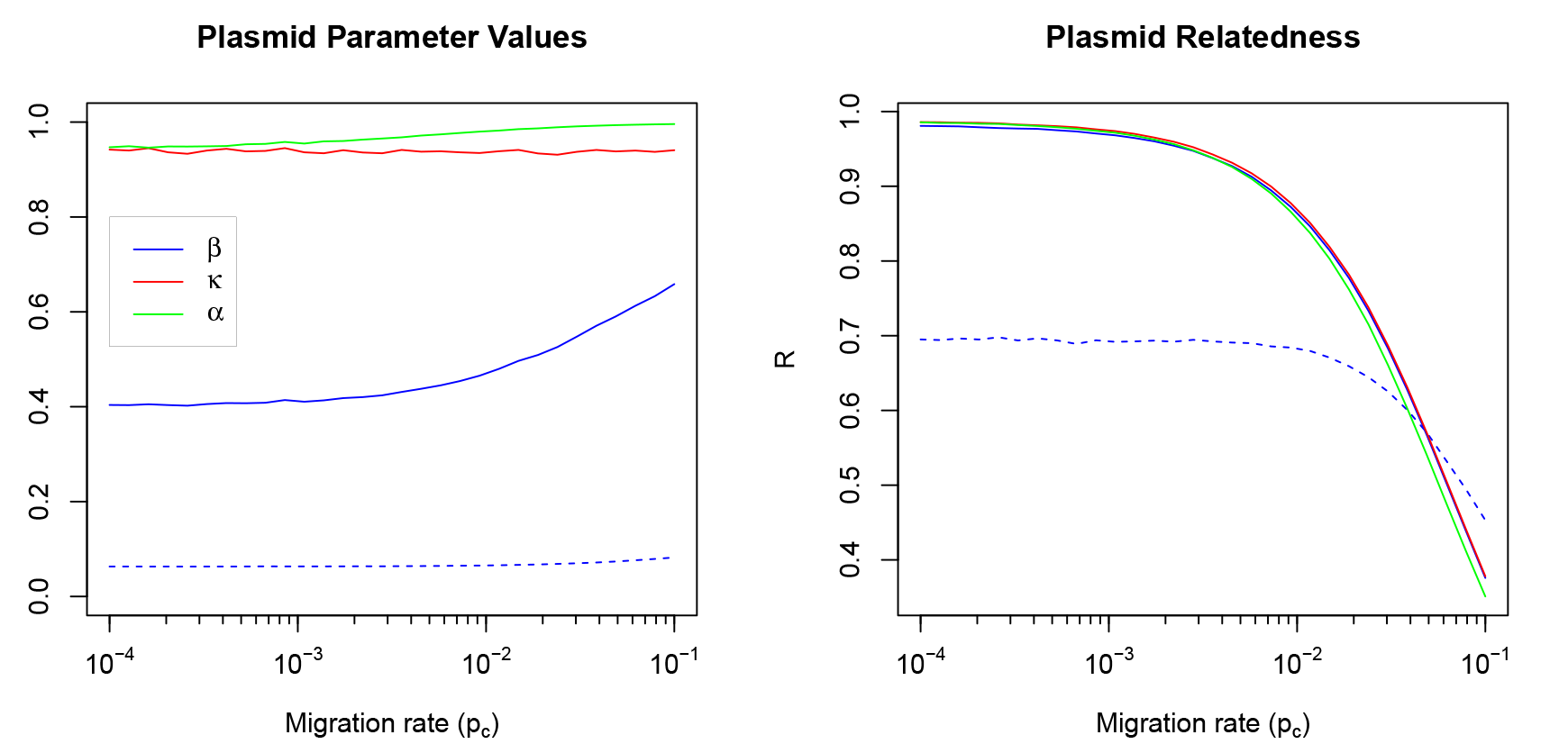
Policing is tuned with declining relatedness. Mean values (left panel) and mean relatedness coefficients (right panel) of the plasmid replication parameters β, κ and α in the presence (CNC - solid lines) and absence (NO-CNC - dashed line) of policing. All values were calculated from the equilibrium phase (second half) of the simulation dynamics, averaged across 100 independent simulations for each value of the migration probability *p_c_*.

The relatedness coefficient with respect to a trait *z* (which can represent either of the plasmid parameters β, κ, or α) is calculated across all plasmids in the population as the regression coefficient of the mean value of the trait 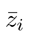 within each host *i* on the trait value *zij* of individual plasmids *j* [28], according to:

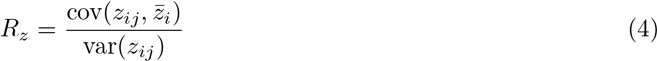

where the calculation of 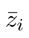 is performed for each focal plasmid by excluding the parameter value of the focal plasmid itself (others-only relatedness). Figure 4 (right) shows that policing has a large positive effect on plasmid relatedness at low and medium migration rates compared to the case where policing is absent. Overall, relatedness declines in response to the increasing mixing of plasmids in the population due to higher levels of migration.

### Migration and selection

The selective pressures on the plasmid replication parameters can be quantified using the Price equation [29,30], a mathematical identity that decomposes the change in the average value of a phenotypic character *z* (e.g. one of the plasmid replication parameters β, κ, or α) into the statistical relationship (covariance) between fitness *f* and the character value *z* among individuals in a population, and the average fitness-weighted errors in the transmission of the character *z* from parent *i* to its offspring (transmission bias), according to:

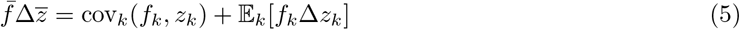
where 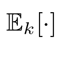 and cov_k_(‧) denote the expectation and covariance over all individuals κ in the population respectively. The Price equation can also be extended to account for individuals that are nested within collectives [30,31], in which case the covariance term representing selection on the phenotypic character z can itself be partitioned into two components, corresponding to the portions of cov(*f*, *z*) that are attributed to selection between hosts (BH) and within hosts (WH) respectively, according to:

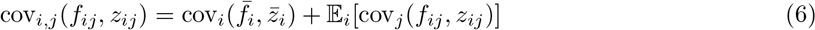
where *i* enumerates hosts, *j* enumerates plasmids in host *i* with plasmid copy number *n_i_*, and 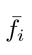 and 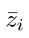 are the average fitness and character values of plasmids in host *i* respectively, with 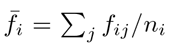 and 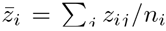. By substituting Equation 6 into Equation 5, we obtain the multi-level version of the Price equation:
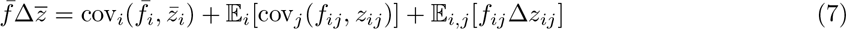

in which the first right-hand-side term captures selection on plasmid collectives at the level of hosts (between-host selection), the second term captures selection on individual plasmids at the intra-cellular level (within-host selection), while the third term quantifies the bias in the transmission of z during plasmid replication.

The components of Equation 7 can be measured in our simulations, thus providing insight into the effects of policing and migration on the interplay between the levels of selection [14]. These components can be further partitioned into the constituent components of plasmid fitness, which is defined as the (actual) number of offspring of plasmid j in the population at the next time step of the simulation. This number includes the plasmid itself (if host i remains alive), any new copies and any possible losses as follows:

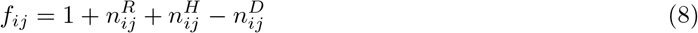
where 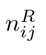 is the number of intra-cellular replication events, 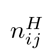 is the number of horizontal transmission events and 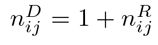 if host *i* dies (randomly or not), otherwise 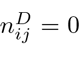. The latter implies that, in the event that host *i* dies, the only copies of plasmid *j* that will be present in the population at the next time step are those that have been transmitted horizontally to another host. This way, between-host selection on character *z* can be expressed as:

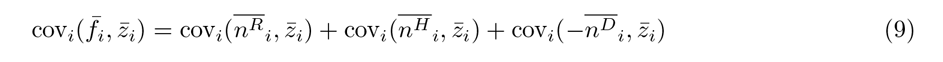
where 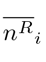 and 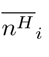 are the average number of events corresponding to replication and horizontal transmission within host *i* respectively, and 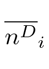 is the average number of plasmid losses due to host deaths.

Similarly, within-host selection on character *z* can be decomposed according to:

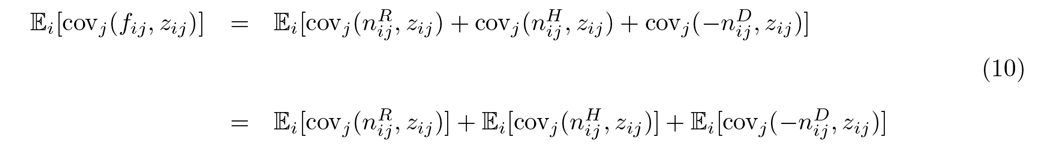

In effect, each component in Equations 9 and 10 captures the portion of between-and within-host selection on character *z* that can be attributed to changes in fitness due to intra-cellular replication, horizontal transmission or losses due to cell death. Figure 5 shows the average values of these components, as well as their sums (the left-hand sides of Equations 9 and 10), across the range of the migration probabilities *p_c_* under consideration. These results demonstrate the progressive intensification of the conflict between the levels of selection across the *p_c_* spectrum for the plasmid replication parameters. Specifically, the selective pressures on the cis-specific (self-regarding) parameters *β* and κ are characterized by the escalating conflict between selection within hosts in favour of selfishness (red solid lines) and selection between hosts in favour of restraint and obedience (blue solid lines). The selective pressures on the trans-acting (other-regarding) parameter *α* are characterized by the contrast between selection between hosts in favour of policing (blue solid line) and the transmission bias (green solid line) which is negative due to the fact that the value of *α* has reached its maximum value at equilibrium, hence mutations that attempt to increase this value further are rejected.

**Figure 5.**
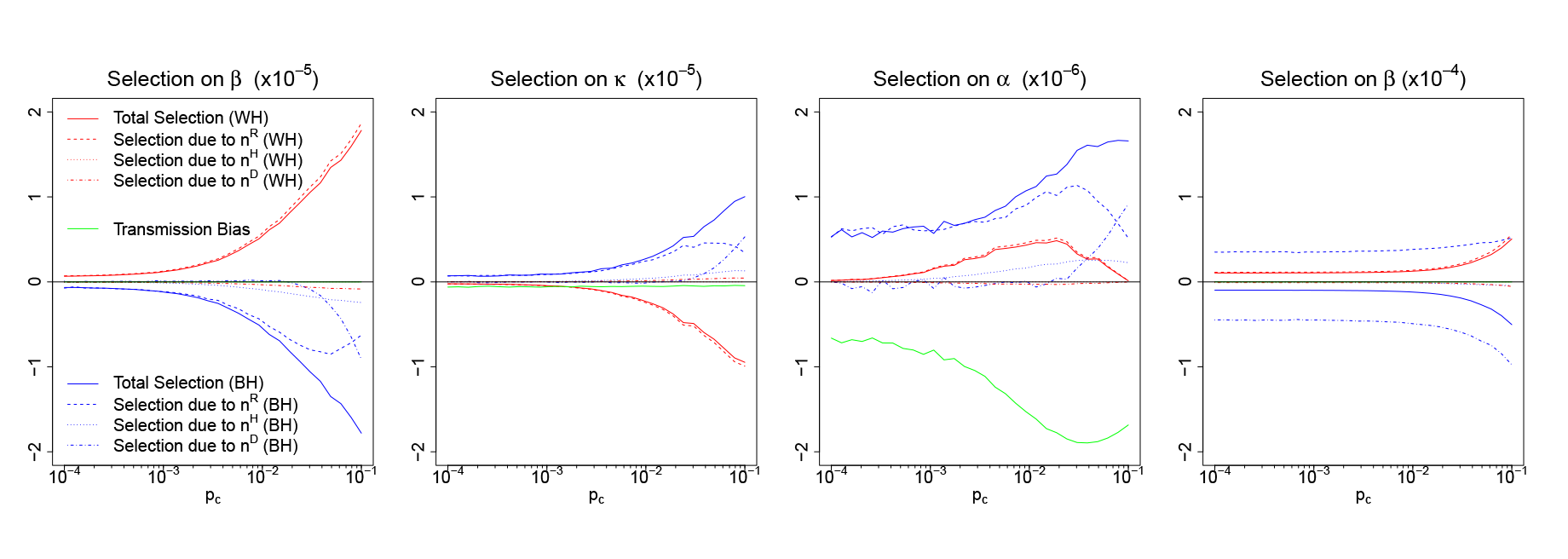
Selective presures on plasmid replication control. The average values of thecomponents of selection on the plasmid replication parameters between hosts (BH; Equation 9) and within hosts (BH; Equation 10), calculated from the equilibrium phase of the simulation dynamics and averaged across 100 independent simulations for each value of the migration probability *p_c_*. Results are provided for simulations that were performed in the presence (5a, 5b, 5c-CNC) and the absence (5d-NO-CNC) of policing.

Figure 5 demonstrates that, at low and moderate migration rates, the dominant factor in shaping selection within and between hosts is intra-cellular replication (blue and red dashed lines). However, the dominance of between-host selection for intra-cellular replication gradually declines (blue dashed lines)at the high migration regime. This indicates a shift in the relative importance of the events that shape selection on the plasmid replication parameters at the level of hosts. Specifically, at low to medium migration rates, it is the slower growth rate of hosts that mitigates the spread of selfish plasmids (blue dashed lines). At high migration rates however, the deterrent of slower host growth is gradually replaced by the risk of host death due to excessive plasmid replication (blue dash-dotted lines). Another factor that makes a moderate contribution in favour of replication control at high migration rates is selection at the level of hosts due to horizontal transmission (blue dotted lines). This reflects the fact that weak replication control in a host (i.e. high 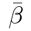 or low 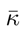) translates into higher copy numbers which lower the per-plasmid horizontal transmission rates, as the probability *p_c_* of horizontal transmission by definition refers to hosts rather than plasmids. This effect is materialized as a positive correlation between replication control and horizontal transmission at the level of hosts. Within-host selection due to horizontal transmission (red dotted lines) is, however, negligible, since the selection of any plasmid for transmission from a donor to a recipient cell is equiprobable.

In the absence of policing (Figure 5d), the selective pressures follow the same general pattern, except for selection at the level of hosts due to intra-cellular replication which actually favours higher basal replication rates (blue dashed line). Positive selection for higher β is compensated by strongly negative selection against β due to host death (blue dash-dotted line), thus yielding a net negative selection against β at the level of hosts (blue solid line). This can be explained by the inherent instability of plasmid replication in the absence of policing, whereby either plasmids will consistently under-replicate, in which case between-host selection will favour higher β, or plasmids will consistently over-replicate in which case between-host selection against β will materialize as resulting from losses due to host death under the weight of excessive plasmid copy numbers [13].

The surge of positive within-host selection on α (Figure 5c - red solid and dashed lines) appears to contradict the intuitive logic that within-host selection on α should be neutral because of the trans-acting nature of replication inhibitors. When a focal plasmid increases its individual production of inhibitors (through higher α), this may or may not increase its growth rate relative to other plasmids within the host, depending on whether it is more or less sensitive to the shared inhibitor in comparison to its neighbours (i.e. whether the focal plasmid’s *κ_ij_* is more or less than the mean 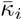 within the host). We would expect the deviation of individual sensitivities from their mean within the host to be, on average, zero, however our results suggest that this is not so for a wide range of *p_c_* values. The key element for interpreting this, seemingly paradoxical, result is shown in Figure 6, which demonstrates a positive relationship between plasmid cis-selfishness (i.e. higher β and/or lower κ) and the production of policing resources (α) in the range of pc values for which we observed the surge of positive within-host selection on α.

**Figure 6.**
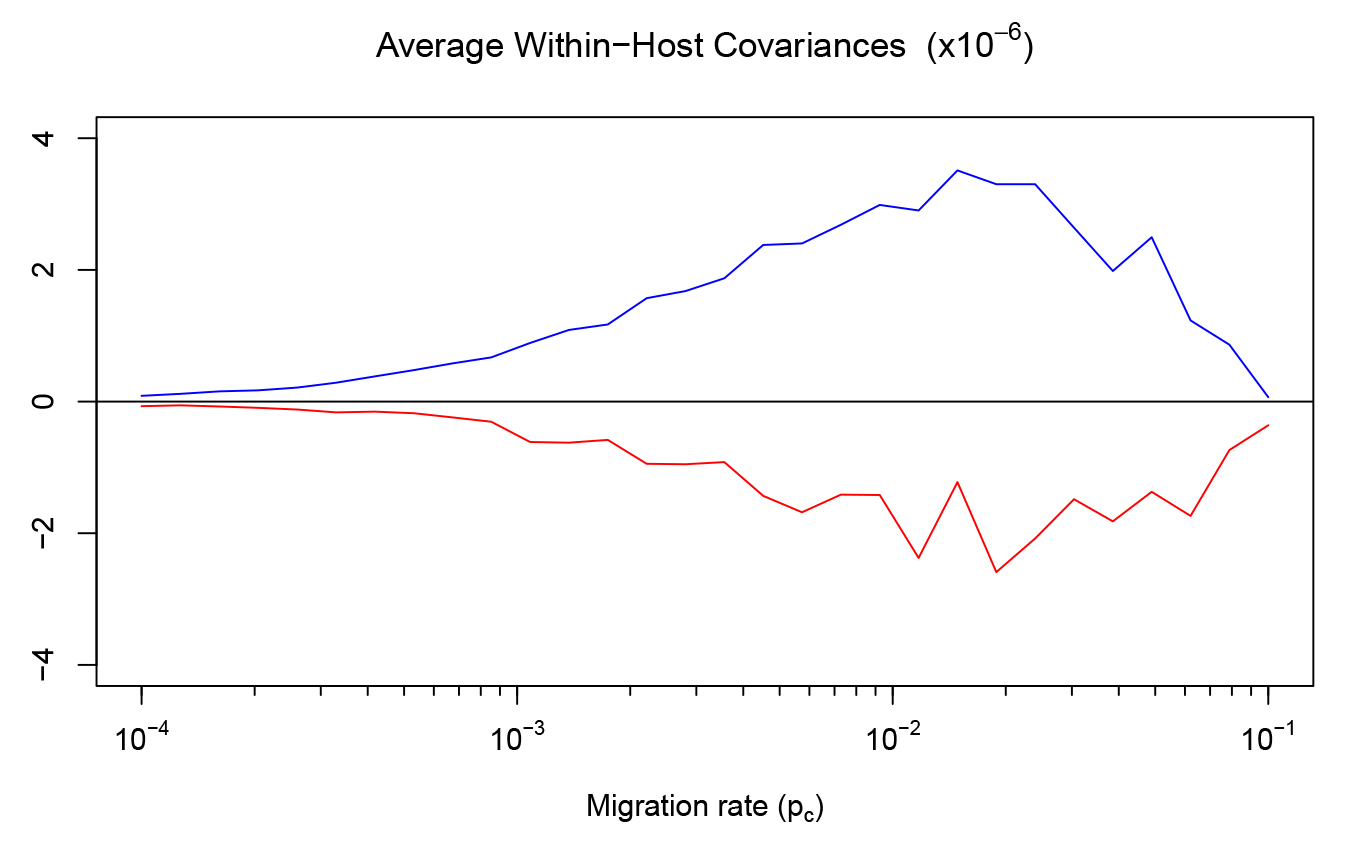
Cis-selfishness mediates the spread of coercive policing. The average values of thewithin-host covariances 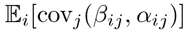 (blue line) and 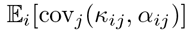 (red line), calculated from the equilibrium phase (second half) of the simulation dynamics and averaged across 100 independent simulations for each value of the migration probability *p_c_*.

## Discussion

Our results illustrate how policing, as a mechanism of repressing competition through collective restraint, offers significant efficiency gains, in terms of host growth and mortality rates, across a wide range of relatedness regimes as determined by varying levels of plasmid migration and mixing across hosts (Figure 1). These efficiency gains were compared to the case where the repression of competition in the plasmid population was achieved through individual restraint in intra-cellular replication. In addition to the performance benefits at the level of hosts, policing also positively contributes towards the improvement of reproductive control and the overall stability of plasmids (Figure 3).

The nature of competition between plasmids in the intra-cellular replication pool is shaped by the CNC mechanism codified by the plasmid replication parameters, namely the cis-acting β (intrinsic selfishness) and κ (sensitivity to policing resources) and the trans-acting α (production of policing resources). These elements of the CNC system are subject to a dynamic evolutionary conflict between two levels of selection [32,33]: within hosts, cis-selfish mutations that increase a plasmid's intrinsic selfishness β or reduce a plasmid’s obedience κ to policing would result in a mutant that would proliferate faster in comparison to more prudent plasmids with a higher degree of adherence to the CNC mechanism. Hence, within-host selection will favour cis-selfishness, thus exacerbating competition between plasmids in the intra-cellular replication pool. However, hosts bearing increasingly selfish and disobedient plasmids will grow slower or even die compared to fellow hosts with stricter replication control among plasmids. This way, within-host selective pressures towards cis-selfishness are counter-balanced by selection at the level of hosts that penalizes hosts with selfish plasmids and, therefore, selfish plasmids themselves.

Our results show that policing evolves across a wide range of mixing regimes and that policing has a strong positive effect on plasmid relatedness at low and medium migration rates compared to the case where plasmids replicate solely on the basis of self-restraint (Figure 4). This positive effect is due to the tighter control of local population size, resulting in more homogeneously sized groups in which individuals are more closely related. Plasmid relatedness decreases with higher migration rates, resulting in a greater degree of coercive policing (higher α).

Using the multi-level form of the Price Equation (Equation 7), we were able to decompose and quantify the selective pressures on the plasmid replication parameters and attribute them to specific classes of events, such as intra-cellular replication, horizontal transmission and death (Equations 9 and 10). Among these, the dominant factor in driving selection in favour of replication control was intracellular replication. At high migration rates, the decline of the portion of between-host selection due to intra-cellular replication, relative to within-host selection, was compensated primarily by the portion of between-host selection attributed to plasmid losses due to cell death. This implies that, at this regime, host death is akin to a sieve that selectively discards the selfish elements that undermine the stability of replication control. At extremely high migration rates (*p_c_* > 0.1), we observed a sharp decline of plasmid obedience to policing leading to the collapse of the population under the weight of excessive plasmid over-replication.

While selection in favour of policing at the level of hosts reflects its positive impact on plasmid collectives with respect to the stability of replication (blue solid line in Figure 5c), the counter-intuitive surge of positive within-host selection on policing (Figure 5c - red solid and dashed lines) can be explained by the positive relationship between plasmid cis-selfishness (i.e. higher α and/or lower κ) and the production of policing resources (α) in the range of *p_c_* values for which we observed the surge of positive within-host selection on α. This indicates that coercive plasmids with higher basal replication rates β and lower obedience κ produce, on average, more policing resources in order to exploit their more prudent neighbours, effectively instructing them to forgo reproduction (through high α) while they continue to replicate themselves (through high β, low κ). When a coercive plasmid is transferred horizontally to a recipient host, in which the resident types are more prudent with respect to replication, the coercive plasmid will have a relative advantage in the local replication pool and will, therefore, leave more offspring. It follows that, since coercive types produce more policing resources on average, this fitness advantage will manifest as positive within-host selection on policing due to intra-cellular replication, i.e. 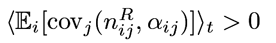(red dashed line in Figure 5c).

We further investigated this hypothesis by setting up additional simulations in which we explored the probability of fixation of a mutant plasmid type in a population dominated by a wild type. Figure 7 demonstrates that the succesful outcome of a mutant invasion depends not only on the mutant’s cis-selfishness but also on its policing trait. Specifically, the successful mutants are more selfish than the wild-type (i.e. have a higher β/lower κ) and, at the same time, produce more policing resources than the wild-type (i.e. higher α), indicating that cis-selfishness mediates the spread of policing which, in turn, facilitates the coevolution of baseline reproduction β and obedience κ towards maximizing the stability of plasmid replication. In the absence of migration, this effect is non-existent as it requires the mixing of selfish and prudent types through among-group migration.

**Figure 7.**
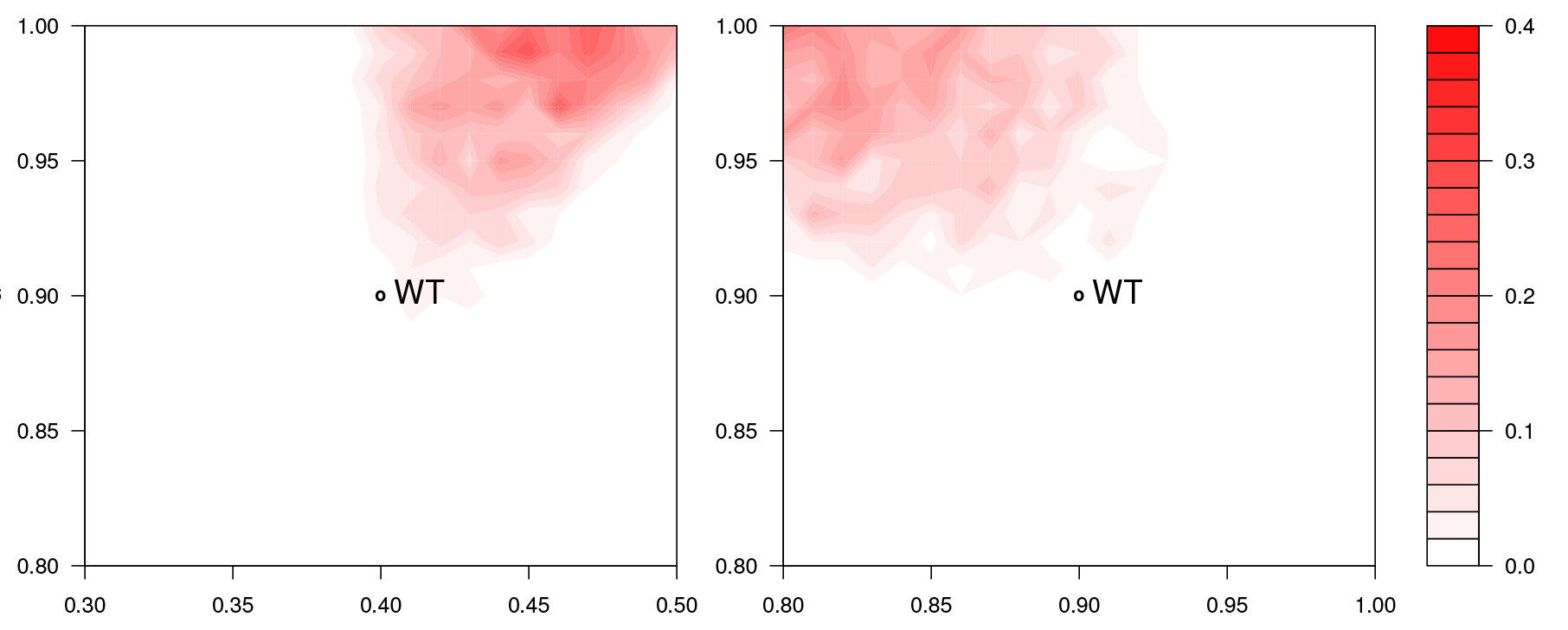
The invasion of cis-selfish mutants. The probability of fixation of a mutant plasmid type in a population dominated by a wild type (WT), measured by exploring the mutant parameter spaces(β, α) with κ = 0.9 (left) and (κ, α) withβ = 0.4 (right) against the same wild type, specified by(β = 0.4, κ = 0.9, α = 0.9). The probability of mutant fixation was calculated by running 100 simulations at each point of the parameter space and counting the number of times the wild-type was replaced by the mutant. Each simulation was seeded with a mixed population of plasmids in which the mutant type measured at 1% of the wild type and the migration probability was set to *p_c_* = 0.01.

We should also note that the enforcement of limits on plasmid parameter values in our simulations (with β, κ, α∈[0,1]), representing various physico-chemical constraints, prevents the continuation of a ratcheting effect in which the succession of selfish and cooperative mutations would be constrained only by the actual costs of producing the initiators and inhibitors of plasmid replication [13]. This effect is present in the coevolution of the plasmid replication parameters within their range of permissible values and stops when α reaches its upper limit. From that point on, the replication control system is stabilized as positive between-host selection on α (which would normally lead to further increases of its value and to corresponding retaliations by the cis-acting β) is counter-balanced by the negative transmission bias (Figure 5c - green line). Overall, the escalating conflict between the cis-selfish and cooperative traits of the plasmid replication system brings increasing stability and control over the local plasmid population size yielding benefits to hosts and plasmids alike.

Finally, the establishment of coercive cheaters under conditions of high mixing is not necessarily limited to a reproductive policing scenario. Coercive strategies are possible whenever there is a joint evolution of a signal trait such as policing (α) and a response trait such as obedience (κ). This opens up the possibility of high signal, low response individuals manipulating low signal, high response individuals. Earlier theoretical work on bacterial communication systems (quorum-sensing) under varying conditions of relatedness highlighted that increased genetic mixing can lead to high signal, low response equilibria [34]. More recently, the phenotypic characterization of signal/response strategies in interacting microbes has uncovered a diversity of signal/response behaviours [35], which is consistent with the existence of coercive signaling strategies.

## Acknowledgments

This work was supported by EPSRC grant EP/H032436/1. KK and RAG are supported by the Medical Research Council, UK.

